# A Novel Partitivirus Confers Dual Contradictory Effect to Its Host Fungus: Growth Attenuation and Virulence Enhancement

**DOI:** 10.1101/2024.09.02.610770

**Authors:** Zhen Liu, Maoqiu Chen, Linghong Kong, Hamdy Aboushedida, Risky Kartika Sari, Hiromitsu Moriyama, Wenxing Xu

## Abstract

Mycoviruses represent potential role in the biocontrol approach due to their ability to reduce both virulence and vegetative growth in some phytopathogenic fungi. However, the mycoviruses enhancing the damage caused by these fungi remains poorly identified and characterized. In this study, a novel double-stranded RNA (dsRNA) fungal virus, tentatively named Sinodiscula camellicola partitivirus 1 (ScPV1), was identified in the phytopathogenic fungus *Sinodiscula camellicola*, isolated from tea leaves. ScPV1 possesses two genomic components of 1835 and 1697 bp, each containing an open reading frame (ORF) that encodes the putative RNA-dependent RNA polymerase (RdRp) and coat protein (CP), respectively, as confirmed by mass spectrometry. Phylogenetic analysis of the amino acid sequences of the RdRps from ScPV1 and related mycoviruses placed ScPV1 within a newly proposed genus, Epsilonpartitivirus, of the family *Partitiviridae*. Viral purification via ultracentrifugation and transmission electron microscopy observations revealed that ScPV1 dsRNA genomes are encapsidated within virus particles approximately 31 nm in size, ranging from 24.9 to 36.8 nm, along with the RdRp protein in an unexpected size. Transfection experiments with purified virions exhibited transfectants with significant reduced growth rates while increased virulence, indicating that ScPV1 has two-sided effect on its host fungus. This finding represents a significant advancement in understanding the complex interactions between mycoviruses and their host fungi.

**Author summary:** To characterize new molecular and biological traits of partitiviruses will provide substantial contributions for a better understanding of mycoviruses, as partitiviruses are prevalent across a wide range of hosts, including plants, protists, and fungi. Here, we identified a novel partitivirus, tentatively named Sinodiscula camellicola partitivirus 1 (ScPV1), from a fungus infecting tea plant, marking the first report of a partitivirus from a phytopathogenic fungus infecting tea plant. ScPV1 is characterized by two dsRNA genomic components encapsidated in particles of varying sizes, along with an RNA-dependent RNA polymerase protein of an expected size and containing some unique amino acids, indicating its distinct molecular and morphological traits. Our biological tests on transfectants generated *via* protoplast transfection with purified virions have rigorously demonstrated that ScPV1 impairs vegetative growth while enhancing virulence in its fungal host. This finding represents the first instance of a mycovirus responsible for hypervirulence and exhibiting dual effects on a phytopathogenic fungus through virion-transfection, as well as the first case of a partitivirus conferring hypervirulence while promoting vegetative growth in a phytopathogenic fungus. We anticipate that these findings will significantly advance our understanding of the complex interactions between mycoviruses and their host fungi.

## Introduction

Mycoviruses (or fungal viruses) either affect the biological traits of their host fungi or keep latent [1]. Some viruses can impair the growth of and reduce the virulence in their host fungi, making them potential biocontrol agents. For example, Cryptonectria hypovirus 1 (CHV1) has been used to control chestnut blight in Europe, and Sclerotinia sclerotiorum hypovirulence-associated DNA virus 1 (SsHADV-1) has been applied to manage rape Sclerotinia stem rot in China [2,3]. Conversely, certain mycoviruses can enhance specific biological traits in their host fungi, granting them an adaptive advantage in particular environments. This is illustrated by the cases of Beauveria bassiana victorivirus 1 (BbVV-1) and Beauveria bassiana polymycovirus 1 (BbPmV-1), which, when infecting the entomopathogenic fungus Beauveria bassiana, lead to an increase in virulence against insects [4]. This hypervirulent trait is very attractive since it enhances the biocontrol effect to the host fungi against agricultural pests. On the other hand, two mycoviruses have been demonstrated to be related to hypervirulence in phytopathogenic fungi: a 6.0-kbp dsRNA, which is phylogenetically related to plant cryptic viruses, in *Nectria radicicola*, the causal fungus of ginseng root rot [5]; and Rosellinia necatrix quadivirus 1 (RnQV1) in *Leptosphaeria biglobosa*, causing *Phoma* stem canker (blackleg) of oilseed rape [6]. The mycoviruses associated with hypervirulence in phytopathogenic fungi have implications that run counter to their potential use as biocontrol agents. Consequently, careful management and utilization of these mycoviruses are essential to prevent them from inadvertently increasing the virulence of their fungal hosts.

The family *Partitiviridae,* with a big member group that respectively infect fungi, plants, and protozoa, has been divided in into five genera, including *Alphapartitivirus*, *Betapartivirus*, *Deltapartivirus*, *Gammapartitivirus*, and *Cryspovirus*, and two newly proposed genera Epsilonpartitivirus and Zetapartitivirus [7–9]. Members of this family possess genomes comprising two linear double-stranded RNA (dsRNA) molecules, sized between 1.4 to 2.4 kilobases (kbp), with the larger molecule encoding the RNA-dependent RNA polymerase (RdRp) and the smaller one encoding the capsid protein (CP), both of which are individually packaged into viral particles measuring 25-40 nanometers in size. [9]. Partitiviruses are generally known for their persistent infections in their hosts [7]; however, there are notable exceptions. For example, Colletotrichum alienum partitivirus 1 (CaPV1) [10] has been shown to significantly reduce host virulence, mycelial growth, appressorial development, and appressorium turgor, while increasing conidial production with abnormal morphology [10]. Similarly, Aspergillus fumigatus partitivirus 1 [11] and Heterobasidion partitivirus 3 [12] have been found to attenuate the growth rates of their host fungi, and Aspergillus flavus partitivirus 1 [8] and Sclerotinia sclerotiorum partitivirus 1 [13] have been observed to attenuate virulence. Additionally, Rosellinia necatrix partitivirus 2 (RnPV2) despite showed no apparent morphological changes to its host fungal *Rosellinia necatrix*, but induced obvious morphological changes in a Dicer-like 2 knockout mutant (dcl-2) of a non-natural host, *Cryphonectria parasitica* causing chestnut blight [14]. Moreover, infection with a partitivirus has been shown to induce hypervirulence in *Talaromyces marneffei*, a thermal dimorphic clinical fungus [15]. However, the extent to which partitivirus-mediated hypervirulence occurs in phytopathogenic fungi in nature is still uncertain.

Tea, represented by the species *Camellia sinensis* (L.) Kuntze, is of considerable economic significance in China and provides a multitude of health benefits to humans. Within the spectrum of diseases that afflict tea plants, anthracnose emerges as one of the most severe, presenting a substantial challenge to the cultivation of tea [16]. The pathogens responsible for tea anthracnose exhibit regional diversity across the globe, with the disease commonly attributed to *Discula theae-sinensis* (formerly known as *Gloeosporium theae-sinensis*) and various species of *Colletotrichum* [17]. Recently, two new fungal species, *Sinodiscula theae-sinensis* and *Sinodiscula camellicola*, which belong to a newly proposed genus Sinodiscula within the family Melanconiellaceae, have been identified and characterized as causative agents of tea anthracnose [18]. The disease caused by these phytopathogenic fungi exhibits similar symptoms: initially, dark green or yellowish-brown watery spots appear, which then expand along the leaf veins, forming irregular-shaped spots. These spots gradually turn brown or reddish-brown and eventually become greyish-white [18]. The visual similarity of the disease symptoms caused by these pathogens presents a challenge in differentiating them through unaided visual inspection. Consequently, a specific disease name, “the tea leaf blight,” has been proposed to describe the symptoms induced by Sinodiscula species [18]. Given that the majority of Chinese tea cultivars are susceptible to this fungal disease, the implementation of effective control measures is essential to protect the yield and quality of tea crops. This imperative necessitates the investigation of mycoviruses that infect the related fungus, as understanding these viral interactions could potentially lead to novel strategies for disease management.

In this study, a novel partitivirus was identified from *S. camellicola* isolated from tea plants, and its molecular, morphological, and biological traits were characterized. This partitivirus exhibits some unique molecular and morphological traits unreported in other partitiviruses, and dual effects on the fungal host by simultaneously attenuating growth while enhancing virulence, marking the first instance of a partitivirus contributing to hypervirulence in a phytopathogenic fungus.

## Materials and methods

### Fungal strains and cultures

*S. camellicola* strain WJT-1-1 was isolated from a diseased tea leaf showing the typical symptoms of leaf blight, while strain SST was isolated from a healthy tea leaf. Both strains were collected from tea gardens in Enshi, Lichuan and Zigui, Hubei Province, China. *S. camellicola* strain WJT-1-1 was found to carry ScPV1 and ScPV1-free *S. camellicola* SST was obtains. The fungal strains were successively cultured on potato dextrose agar (PDA; 20% diced potatoes, 2% glucose, and 1.5% agar) medium at room temperature (∼25°C) unless otherwise stated.

### Extraction and validation of strain dsRNA

For dsRNA extraction, mycelial plugs were inoculated onto sterilized cellophane disks on PDA plates for 3 to 5 days. Frozen mycelial powder was suspended in SDS buffer. The column method was carried out to extract total RNA of the strain as previously described [19] and eluted with RNase-free water. Residual DNA and ssRNA were eliminated from the extracted nucleic acids using 2U DNase I (Simgen, Hangzhou, China) and 10U S1 nuclease (TaKaRa, Dalian, China) at 37°C for 1 h. Purified dsRNA were subjected to electrophoretic separation on a 1.2% (w/v) agarose gel utilizing Tris-acetate-EDTA (TAE) buffer, and the resulting fractions were subsequently visualized through staining with ethidium bromide. ScPV1 genomic dsRNAs was excised, purified using DNA Gel Extraction Kit (Simgen, Hangzhou, China), dissolved in RNase-free water and stored at −70 °C until use.

### Full-Length Genome Amplification and Sequencing

The sequences of the two genomic dsRNAs of ScPV1 were determined by cloning and sequencing processes involving the generation of amplicons by RT-PCR employing the random primers 05RACE-3RT and 05RACE-3 (S2 Table), as previously described [20]. The terminal sequences at both the 5′ and 3′ ends of the dsRNAs were acquired through cloning and sequencing of the RT-PCR amplicons produced via a standardized RNA ligase-mediated rapid amplification of cDNA ends (RLM-RACE) protocol, as detailed in S2 Table. The oligonucleotide primers utilized in the RLM-RACE process were synthesized based on the sequence data procured from the randomly primed amplicons [21]. The recombinant clones were sequenced by Sanger sequencing (Sangon Biotech Company, Ltd., Shanghai, China) using the universal primer pair M13F-47/M13R-48. At least three independent clones were sequenced.

### Sequence Assembly and Phylogenetic Analysis

Sequences were analyzed and assembled with SnapGene software. The obtained full-length sequences were searched for sequences with homology comparison in the BLAST programme of the NCBI (http://www.ncbi.nlm.nih.gov) database. ORFfinder (https://www.ncbi.nlm.nih.gov/orffinder/) was used to predict possible open reading frames. Nucleic acid sequences and amino acid sequences were multiplexed using the software MAFFT (https://www.ebi.ac.uk/jdispatcher/msa/mafft) and use GeneDoc to visualize the results. Phylogenetic tree analysis was performed using MEGA 7 software based on the coding virus RdRp nucleotide sequence using the neighbour-joining method [22]. Self-expansion support of the generated phylogenetic trees was examined with the Bootstrap program (1000 replicates). The online websites Phyre2 (http://www.sbg.bio.ic.ac.uk/phyre2/) and SMART (http://smart.embl-heidelberg.de/) to predict protein conserved structural domains.

### Purification analysis and electron microscopic observation of virus particles

Extraction of ScPV1 virus particles from strain WJT-1-1 and negative control strain SST-5 was performed using ultrafast gradient centrifugation as previously described [23]. Briefly, *ca.* 30 g mycelia were mixed with 100 mM phosphate buffer (PB; 8.0 mM Na_2_HPO_4_,2.0mM NaH_2_PO_4_, pH 7.4) at a ratio of 4 mL/g of mycelia, followed by centrifugation at 10000 rpm/min for 30 minutes at 4 °C to eliminate cellular debris. Subsequently, the supernatant was subjected to ultracentrifugation (utilizing the Optima LE-80K system from Beckman Coulter, Inc.) at 26000 rpm for 3 hours at 4 °C, enabling the precipitation of viral particles. These particles were then resuspended in a 100 mM phosphate buffer. Further purification of the crude viral preparation was achieved through sucrose gradient centrifugation [24]. Subsequently, aliquots of 100 μL from each fraction underwent double-stranded RNA (dsRNA) extraction to ascertain the presence of viral dsRNAs. The purified virion solution was adsorbed onto a copper mesh for 3 minutes and then the mesh was placed on absorbent filter paper and allowed to dry for 2 minutes. Finally, the mesh was placed in a negative staining solution for 3 minutes and dried for 2 minutes for electron microscopic observation [25]. The length of the observed viruses was measured using ImageJ software [26].

### SDS-polyacrylamide gel electrophoresis

Proteins isolated from distinct sucrose gradient fractions underwent comprehensive fractionation procedures utilizing 12% SDS-polyacrylamide gel electrophoresis (PAGE). This electrophoretic analysis was performed in a buffer system composed of 25 mM Tris/glycine and 0.1% SDS, which provided optimal conditions for protein separation. The resulting gels were subjected to staining with Coomassie brilliant blue R-250 (Bio-Safe CBB; Bio-Rad, USA), and individual protein bands were excised for peptide mass fingerprinting (PMF) analysis by Sangon Biotech, Co., Ltd, in Shanghai, China [27].

### Analysis of dsRNAs on horizontal transmission

Confrontation culture method was used to determine the horizontal infectivity of dsRNA viruses [28]. After culturing the WJT-1-1 (ScPV1-infected; donor) and SST-5 (ScPV1-free; recipient) strains on PDA plates, mycelial clumps were punched out from the edge of the colonies with a sterilized punch (5 mm diameter). The mycelial discs of strains WJT-1-1 and SST-5 were co-cultured together in 9 cm diameter Petri dishes at 25°C, with a total of six replicates. After seven days of incubation, mycelial clusters were collected from the peripheral hyphal growth area of strain SST-5 and sub-cultured onto PDA plates covered with sterilized cellophane. A total of 24 mycelial discs were selected, and dsRNAs were extracted and detected following an additional seven days of incubation at 25°C in a constant temperature incubator.

### Biological testing

Biological measurements consist mainly of growth rate and pathogenicity measurements [29]. The tested strains were inoculated onto PDA plates and incubated in the darkness at 25°C. When the mycelium grew to about 3 cm, a number of actively growing mycelium blocks were punched at the edge of the colony with a hole punch (5 mm in diameter) inside a sterile laminar air flow, transferred to the centre of new PDA plates, and cultured in a constant temperature incubator at 25°C, with three replicates for each strain. Ten days post inoculation, the colony diameter was measured by the criss-cross method, the data were recorded, and the average growth rate was calculated. Virulence tests were conducted on fresh detached tea leaves (*C. sinensis* cv. ‘Echa 1’) in six replicates. Briefly, tea leaves were subjected to a three-step washing process with sterile water, followed by air-drying, prior to being inoculated. The leaves were pricked three times with a sterilized needle (0.5 mm in diameter) and inoculated with a mycelial disc. The inoculated leaves were incubated at 25°C and disease progress was recorded every 24 h.

### Preparation of Protoplasts and Transfection of Virus Particles

Protoplasts were prepared from fresh mycelium of the ScPV1-free *S. camellicola* strain SST-5 that were growing vigorously as described previously [30]. Briefly, *ca.* 0.5 g mycelia were mixed with 10 mL NaCl buffer (700 mM NaCl, 0.1 g lysing enzymes from trichodema harzianum and 0.01 g snailase), followed by cultivation at 100 rpm/min for 3 h at 30°C to obtain protoplasts. Subsequently, protoplasts were filtered using a Millipore filter and counted with a hemocytometer slide, 2.0×10^6^ protoplasts were used for transfection with *ca.* 70.0–80.0 μg ScPV1 virions were transfected with PEG 6000 as previously described [31]. After transfection, the protoplast suspension was evenly spread on Bottom Agar plates (200 g of sucrose, 3 g of yeast extract per liter 3 g casein acid hydrolysate, 10 g agar) and incubated at 25°C.

### Statistical analysis

The biological data were statistical analysis using SPSS Statistics Software 27.0 with one-way analysis of variance (ANOVA) and comparison of means. The mean values for the biological replicates are presented as column charts with error bars representing standard error of mean (SEM). The graphs were generated using both MS Excel and GraphPad Prism 7 (a software program provided by GraphPad). *P*-values < 0.05 were considered to indicate statistical significance.

## Results

### Patitiviral dsRNAs were detected in *S. camellicola* strain WJT-1-1

A total of nucleic acids were extracted from the mycelia of *S. camellicola* strain WJT-1-1 and strain SST-5 after digestion with DNase I and S1 nuclease, as illustrated using agarose gel electrophoresis, it reveals two dsRNA bands (termed dsRNA1 and 2 according to their decreased sizes) exclusively in strain WJT-1-1 (Fig 1A and 1B). Following random primers for reverse transcription and polymerase chain reaction (RT-PCR) and rapid amplification of cDNA ends (RACE), the sequences of full-length complementary DNA (cDNA) clones were determined for dsRNAs 1 (GenBank No. PQ201539) with 1835 bp and dsRNA2 (PQ201540) with 1697 bp (Fig 1C). Genomic organization analysis of both dsRNAs revealed that dsRNA1 contains one open reading frame (ORF) in positive strand which starts at position 79 nt and ends at position 1788 nt with a stop codon encoding a putative RdRp of 569 amino acids (aa) and an estimated molecular weight of 65.5 kDa. The ORF2 spanning from position 101 nt to 1606 nt in dsRNA2 encoded a putative CP of 501 aa with molecular weight of 55.7 kDa.

**Fig. 1.**
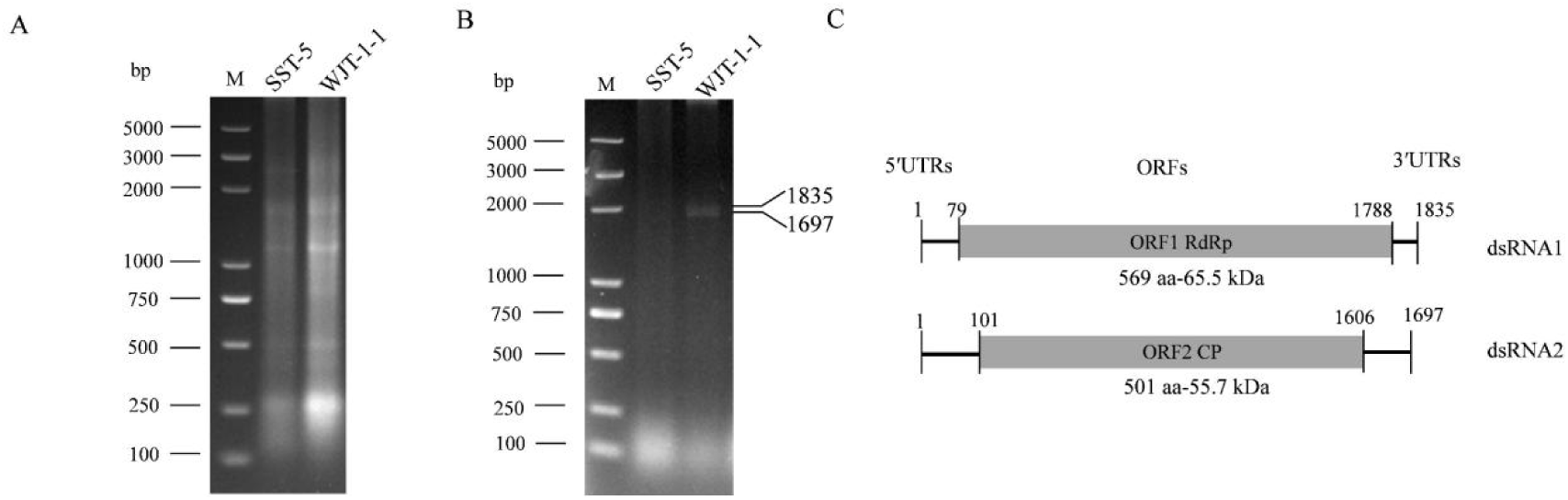
Electrophoresis analysis and genomic organization of Sinodiscula camellicola partitivirus1 (ScPV1). (A and B) Electrophoretic profile on agarose gels of dsRNA preparations extracted from *Sinodiscula camellicola* strains SST-5 and WJT-1-1 before (A) and after treatment with DNase I and S1 nuclease (B), respectively. (C) Schematic representation of the genomic organization of ScPV1 RNA 1 and RNA 2. Grey boxes and black lines indicate open reading frames (ORFs) and untranslated terminal regions (UTRs), respectively.

Blastx searches revealed that the proteins encoded by dsRNA1 and dsRNA2 shared the highest identity with RdRp (NCBI GenBank No. UOK20169, 92% coverage, 62.70% identity, E value = 0.0) and CP (88% coverage, 52.27% identity, E value = 5e-160) of Diplodia seriata partitivirus 1 (DsPV1) and Colletotrichum liriopes partitivirus 1 (ClPV1) (S1 Table), supporting that dsRNAs 1 and 2 harbor a putative ORF encoding RdRp and CP, respectively. Additionally, putative RdRp shared serially high identifies (92%-93% coverage, 61.71%-62.46% identity, E value = 0.0) with RdRp of Colletotrichum eremochloae partitivirus 1, Colletotrichum liriopes partitivirus 1, Erysiphe necator associated partiti-like virus 1 and Metarhizium brunneum partitivirus 1 in the family *Partitiviridae* (S1 Table). Based on their molecular characteristics and similarity to other viruses, these dsRNAs are proposed as a novel partitivirus, tentatively named Sinodiscula camellicola partitivirus 1 (ScPV1).

### Multiple alignment and phylogenetic analysis of the RdRp reveals that ScPV1 belongs to a new viral genus

Multiple alignment analysis was performed for the putative RdRp sequences of ScPV1 and some closely related members in the same family. Six conserved motifs (Motifs Ⅲ to Ⅷ), similar to those observed in other related members, were detected, whereas ScPV1 RdRp indicated several unique amino aicds, including a phenylanaline (F) instead of leucine (L) in Motif IV, F instead of Valine (V) or Isoleucine (I) or L in Motif V, and Serine (S) instead of aspartic acid (D) in Motif VII (Fig 2A). A phylogenetic tree was constructed using the RNA-dependent RNA polymerases (RdRps) of ScPV1 and representative members from various genera within the family Partitiviridae. The analysis reveals that the RdRp of ScPV1 clusters with those of members belonging to the newly proposed genus Epsilonpartitivirus, while also forming an independent branch within the tree (Fig 2B). Based on the established threshold criteria for species distinction, which requires a 90% similarity in RdRp sequences [32], ScPV1 is identified as a novel member of the genus Epsilonpartitivirus within the family *Partitiviridae*.

**Fig. 2.**
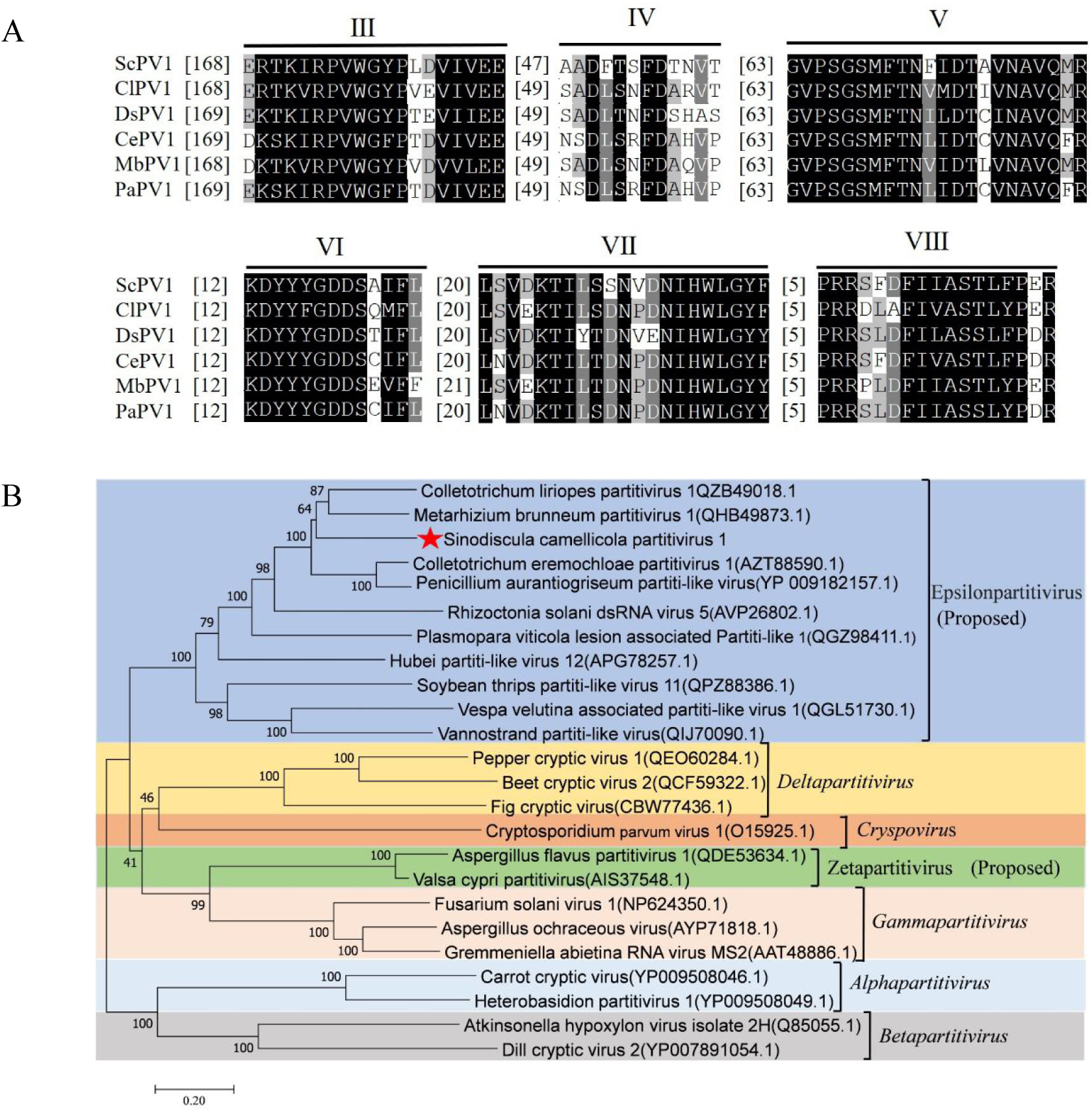
Sequence and phylogenetic analysis of ScPV1 genome. (A) Multiple alignment of six conserved motifs (III to VIII) in the RNA-dependent RNA polymerase (RdRp) sequences of ScPV1 and some close related members belonging to the family *Partitiviridae.* Black and gray backgrounds indicated conserved and semi-conserved amino acid residues, respectively. The abbreviated names refer to Colletotrichum liriopes partitivirus 1(CIPV1, QZB49018.1); Colletotrichum eremochloae partitivirus 1(CePV1, AZT88590.1); Metarhizium brunneum partitivirus 1(MbPV1, QHB49873.1); Diplodia seriata partitivirus 1(DsPV1, UOK20169.1); and Penicillium aurantiogriseum partitivirus 1(PaPV1, AZT88600.1). (B) A maximum likelihood phylogenetic tree constructed using the the RdRp sequences of ScPV1 and some representative members of all the genera belonging to the family *Partitiviridae*. The red pentagram indicates the position of ScPV1.

### ScPV1 were encapsidated in isometric particles

To determine the viral particles of ScPV1, assumed virus particles were purified from strains WJT-1-1 and SST-5, and subjected to stepwise sucrose gradient centrifugation (10% to 40% with 10% sucrose increments). Nucleic acids were extracted from each sucrose gradient fraction and loaded for electrophoresis analysis on 6% PAGE gel, and the results showed that the target dsRNA bands, matching the sizes of those extracted from the mycelia of strain WJT-1-1, were exclusively detected in the nucleic acid preparation of strain WJT-1-1, sharply concentrated in the 30% sucrose fraction (Fig 3A).

**Fig. 3.**
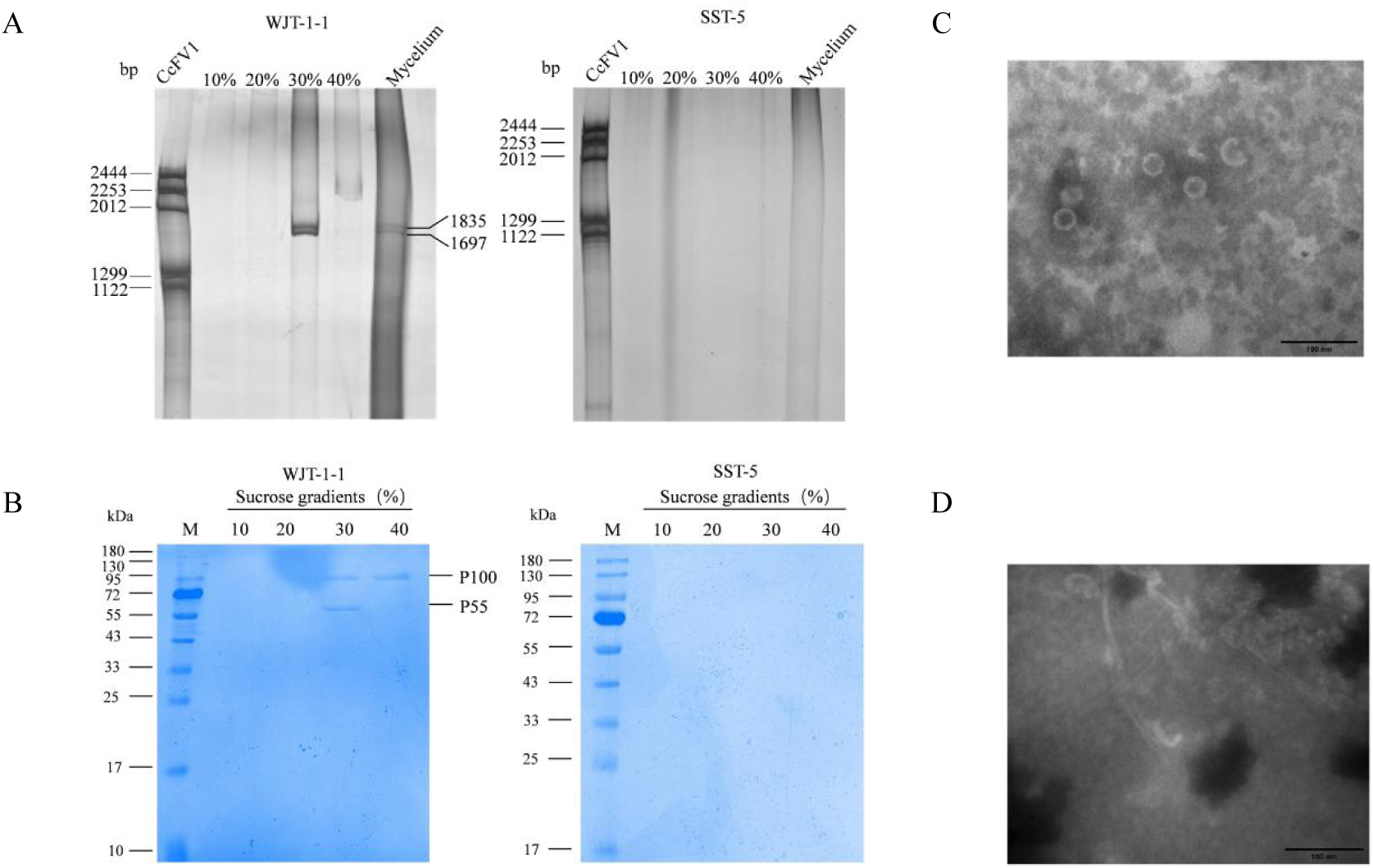
Analysis of nucleic acids and proteins associated with ScPV1 virus-like particles (VLPs). (A) Agarose gel electrophoresis of dsRNAs extracted from purified ScPV1 VLPs obtained from sucrose gradient fractions ranging from 100 to 400 mg/mL with an increment of 100 mg/mL, and from the mycelia of strains WJT-1-1. Colletotrichum camelliae filamentous virus 1 (CcFV-1) genomic components were involved as a dsRNA marker. (B) SDS-PAGE of proteins extracted from the aforementioned sucrose gradient fractions. M, protein molecular weight marker. (C) Representative TEM images of VLPs purified from strain WJT-1-1. (D) Representative TEM images of VLPs purified from strain SST-5.

To identify the structural proteins, assumed proteins were extracted from the 10% to 40% sucrose gradient fractions and respectively subjected to SDS-PAGE electrophoresis. The results indicated that two protein bands, designated as P55 and P100 according to their estimated sizes, were predominantly found in the 30% sucrose fraction. Additionally, a protein band matching the size of P100 was specifically concentrated in the 40% sucrose fraction. Both protein bands were excised and analyzed using peptide mass fingerprinting (PMF). The results revealed that 228 peptide segments from P55 matched the protein sequence encoded by ORF2, representing approximately 74% of the entire sequence (S3 Table), confirming that P55 is the CP encoded by dsRNA2. Additionally, seven peptide segments from P100 matched the protein sequence encoded by ORF1, accounting for about 20% of the entire sequence (S4 Table), indicating that P100 is the RdRp protein encoded by dsRNA1. In contrast, no corresponding protein bands extracted from ScPV1-free strain SST-5 were detected in any sucrose layer (Fig 3B).

These investigations revealed that ScPV1 dsRNAs co-purified with two protein bands exhibiting molecular weights of 55 and 100 kDa, attaining the highest concentration in the 30% sucrose fraction (depicted in Fig 3B). Transmission electron microscopy (TEM) examination of these fractions revealed the presence of isometric virus-like particles (VLPs), having a diameter that spanned from 24.9 to 36.8 nm, with an average of 31 nm (Fig 3C). In contrast, no viral particles were discerned in the control strain SST-5 (Fig 3D).

### ScPV1 infection reduces the growth rate while increases the virulence of the host fungus

To assess viral transmission, the ScPV1-infected strain WJT-1-1, acting as the donor of the virus, was cultured in direct contact with the uninfected strain SST-5, which served as the recipient. Following a 7-day contact-culture period, 24 mycelial discs (marked by asterisks in Fig 4A) were excised from the colonies of strain SST-5 at six independent peripheral positions. These discs were then subjected to dsRNA extraction for further analysis. After treatment with S1 enzyme, the extracted dsRNA was visualized on electrophoretic agarose gel, and results showed that no ScPV1 dsRNAs were detected in the subisolates of strain SST-5, suggesting that ScPV1 is most likely difficult to be horizontally transmitted from strain WJT-1-1 to another strain (Fig 4B).

**Fig. 4.**
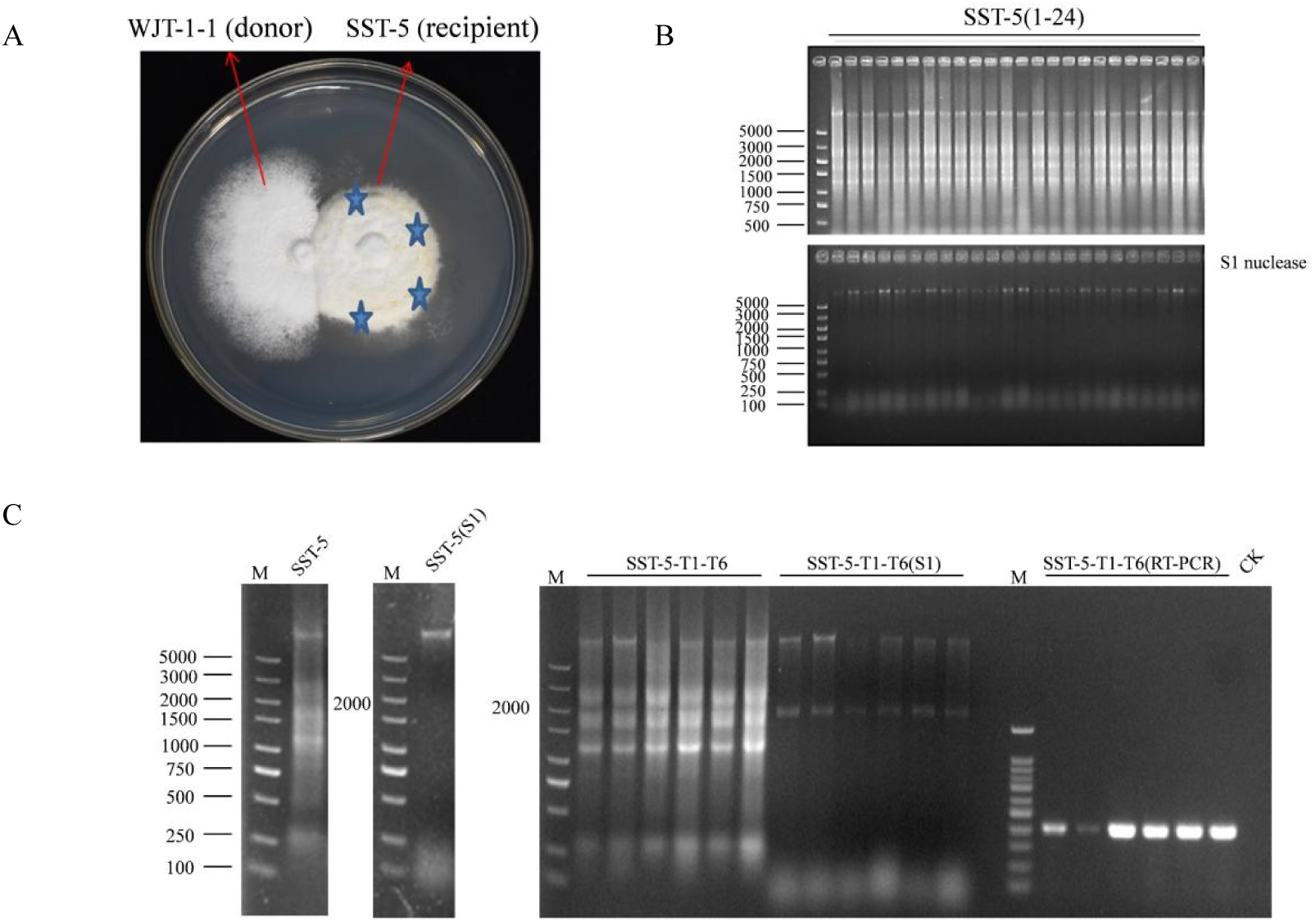
The horizontal transmission effect of ScPV1 and the transfection of ScPV1 virus particles. (A) The representative colony morphologies of strains WJT-1-1 and SST-5 in contact culture. (B) Agarose gel electrophoresis of nucleic acids extracted from strain SST-5 mycelia before (upper panel) and after treatment with S1 nuclease (bottom). (C) Nucleic acids extracted from the transfectants before (left) and after treatment with S1 nuclease (middle), and RT-PCR identification (right) of ScPV1 in the transfectants after transfection with purified ScPV1 VLPs, with the parent strain SST-5 (CK^−^) involved as a negative control.

To further assess the biological traits of the virus, the purified virus particles of ScPV1 were transfected into the protoplasts of strain SST-5 using PEG-mediated methods. A total of 110 newly protoplast-generated colonies were randomly picked up, transferred into new plates for growth, and subjected to nucleic acid extraction. The extracted nucleic acids were treated with S1 enzyme and analyzed via agarose gel electrophoresis, which revealed that six transfectants (representing a frequency of 5%) were infected with ScPV1. This finding was further supported by RT-PCR identification using the extracted nucleic acids and the primer pair named RNA2-F2/RNA2-R2 (Fig 4C and S2 Table).

As cultured on potato dextrose agar (PDA) medium, all the transfectants exhibited sparse hyphal patterns with petal shaped edges, similar to the morphologies of strain WJT-1-1 (Fig 5A). Furthermore, all the transfectants demonstrated reduced growth rates, ranging from 5.9 to 6.3 mm/day, compared to their parent strain SST-5, which exhibited a growth rate of 6.5 mm/day. In contrast, all the transfectants exhibited increased virulence levels, resulting in lesions with lengths of 12.3 to 12.8 mm, compared to lesions (10.9-11.2 mm) caused by strain SST-5 at 7 days post inoculation (dpi) on tea leaves of the cultivar ‘Echa 1’ of *Camellia sinensis* (Fig 5B to 5D).

**Fig. 5.**
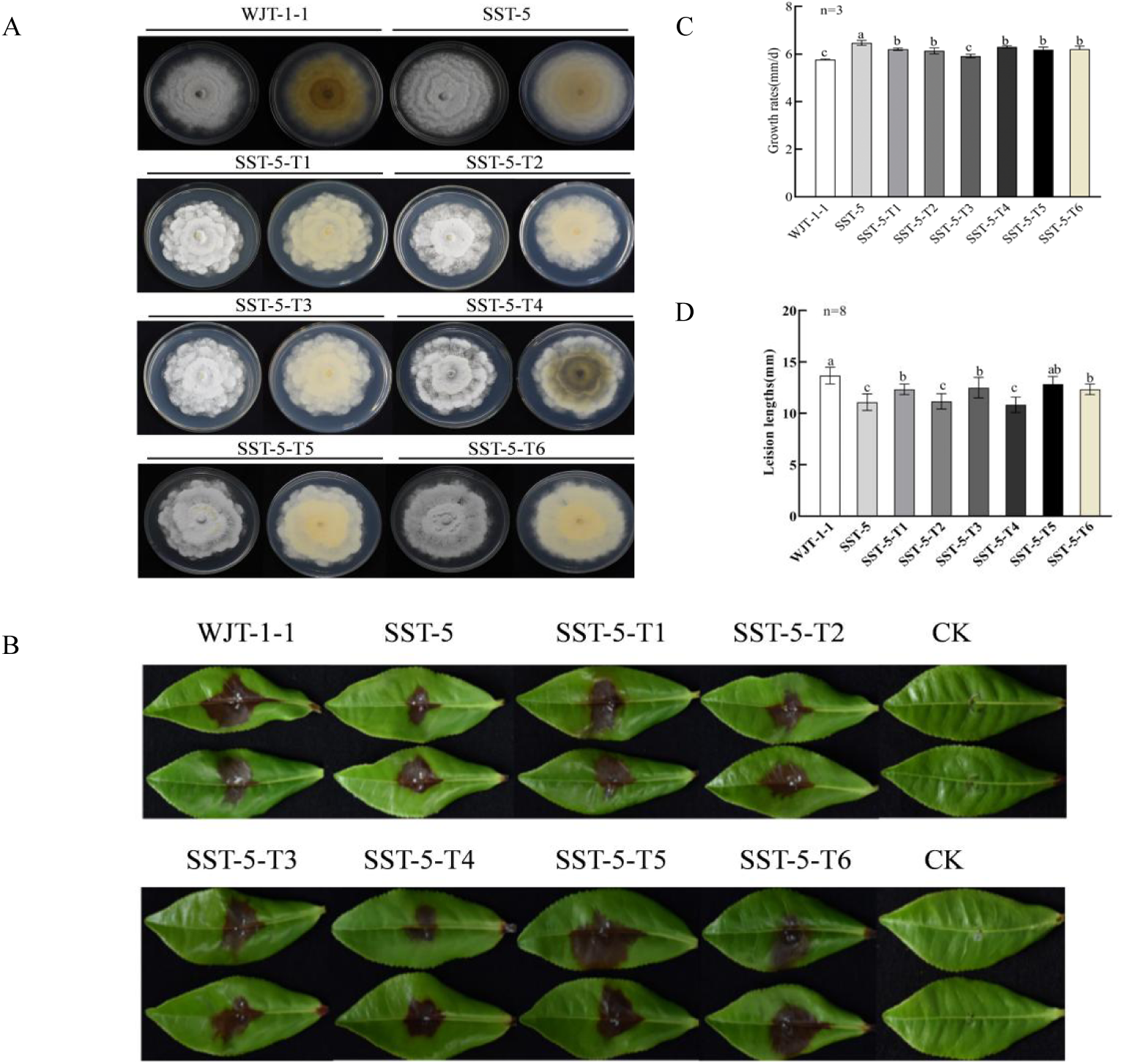
Effects of ScPV1 on fungal morphology, growth, and pathogenicity. (A) Colony morphologies of strain SST-5 and its transfectants (SST-5-T1 to -T6) infected by ScPV1, together with strain WJT-1-1 grown on PDA media at 28 °C in darkness for 5 days. (B) Representative symptoms on tea leaves (*Camellia sinensis* cv. ‘Echa 1’) following inoculation with the aforementioned ScPV1-infected transfectants, strain WJT-1-1, and ScPV1-free ScPV1 at 7 dpi. (C and D) Growth rates (C) and the resulting lesion lengths (D) of the aforementioned strains; columns indicate the average values of independent repeats (n), the different letters refer to significant difference, and error bars represent standard deviation.

## Discussion

In this study, a new mycovirus, tentatively named ScPV1, were identified and characterized from *S. camellicola* isolated from tea plants. According to the species demarcation criteria within the family *Partitiviridae* (less than 90% aa sequence identity in the RdRP [32], as well as its genomic organization and phylogenetic analysis, ScPV1 was determined as a new member belonging to the newly proposed genus Epsilonpartitivirus of the family *Partitiviridae*. To date, five mycoviruses, namely Colletotrichum camelliae filamentous virus 1 (CcFV1, belonging to the family *Polymycoviridae*), Pestalotiopsis theae chrysovirus 1 (PtCV1, *Chrysoviridae*), Pestalotiopsis fici hypovirus 1 (PfHV1, *Hypoviridae*), Melanconiella theae mitovirus 1 (MtMV1, *Mitoviridae*), Didymella theifolia botybirnavirus 1 (DtBRV1, *Botybirnaviridae*) have been identified from phytopathogenic fungi infecting tea plants, including *Colletotrichum camelliae*, *Pestalotiopsis theae*, *Pestalotiopsis fic*i, *S. camellicola,* and *Didymella theifolia*, respectively; while no partitiviruses had been reported in the fungi infecting tea plants [33–37]. To our knowledge, this is the first report of a partitivirus from a phytopathogenic fungus infecting tea plant.

The genomic sequences of two dsRNA fragments of ScPV1 were determined, and their sizes were found to be 1835 bp and 1697 bp, respectively, and conforms to the genomic size range of the family *Partitiviridae,* whose members typically have 1 or 2 segments of 1.4–2.3 kbp per segment [38]. Each dsRNA of ScPV1 is concluded to contain one single ORF, encoding RdRp and CP in dsRNA1 and dsRNA2, respectively, a common feature within *Partitiviridae* family [39]. To investigate the virus particles of ScPV1, assumed proteins were extracted from strain WJT-1-1 and SST-5, and a kind of icosahedral virus-like particles (diameters, approximately 31 nm) were exclusively purified and visualized *via* TEM from the mycelia of strain WJT-1-1 infected by ScPV1 (Fig 3C). Additionally, dsRNAs 1 and 2 were sharply concentrated in 30% sucrose fraction where both virus-like particles and structural proteins were detected (Fig 4C). Additionally, P55, which corresponds in size to the capsid protein (CP) encoded by ORF2 of ScPV1, was identified through SDS-PAGE and peptide mass fingerprinting (PMF) analysis (refer to S3 Table in the supplemental material) (Fig 3B). Altogether these results support the suggestion that dsRNAs 1 and 2 of ScPV1 are encapsidated in the 31-nm viral particles constituted by P55. A major structural protein encoded by ScPV1 dsRNA 2 suggests an icosahedral T=1 capsid consisting of 60 CP polypeptide, which matches the previous partitiviral particle structures such as Fusarium poae virus 1 and Penicillium stoloniferum viruses S and F, characterized by cryo-electron microscopy (cryo-EM), three-dimensional image reconstruction, and X-ray crystallography [40–42]. TEM observation demonstrated that ScPV1 forms isometric particles 24.9–36.8 nm in diameter, fitting in the size range (25–40 nm) of the known members of the families *Partitiviridae* [10], but has more divergent sizes.

It is worthy to note that a P100 protein was detected on SDS-PAGE, and PMF analysis revealed that P100 generated seven peptide fragment that matched the deduced ScPV1 RdRp sequence at amino acid position 64–227, accounting for 20% of the entire coverage. This suggests that P100 is most likely the viral RdRp, which was significantly larger than the predicted MW of 65.5 kDa, possibly due to some processing modifications. Notably, RdRp was concentrated at a similar titer in the 30% sucrose fraction while at a significantly higher level in the 40% sucrose fraction as compared with the concentration of CP bands on the SDS-PAGE gel (Fig 3B), a pattern not observed in other known mycoviruses. Typically, one or two molecules of RdRp are packaged within each particle of a partitivirus [9], so the peptide concentration ratio should be approximately 60:1 or 30:1 for CP versus RdRp in a partitivirus. Multiple alignments of RdRp sequences of ScPV1 and related dsRNA viruses indicate six conserved motifs, namely motifs III to VIII, and the triplet “GDD” in Motif-VI, which is conserved in the RdRp sequences accross +ssRNA and most dsRNA viruses (Fig 2A) [43], while some unique amino acids within Motifs IV, V, and VII were observed for ScPV1 RdRp.

Fungal partitiviruses are transmitted intracellularly during cell division, hyphal anastomosis and sporogenesis, and alpha- and betapartitiviruses infecting fungal hosts from the genus *Heterobasidion* are transmitted *via* hyphal contacts between somatically incompatible host strains [7]. Whereas, horizontal transmission experiments were performed with two different strains WJT-1-1 and SST-5 without success, which might be due to vegetative incompatibility between the donor and the recipient strains. To date, experimental transfection of virions into protoplasts for partitiviruses has been reported only in a few fungal species, including *R. necatrix*, *Aspergillus fumigatus*, and *S. sclerotiorum* [11,13,14,44]. In this study, vertical transmission was achieved using purified particles, and ScPV1 was shown to be transmitted at a low frequency of 5%, which facilitates a critical assessment of the biological traits of ScPV1.

Most of the viruses in the family *Partitiviridae* are latent infection in their host fungi, but some of them can attenuate the fungal virulence and other biological activities, thus are considered as potential biological agents. For example, Colletotrichum liriopes partitivirus 1 (ClPV1) could reduce the virulence and sporulation of *Colletotrichum liriopes* [45]; Colletotrichum alienum partitivirus 1 (CaPV1) significantly decreased host virulence, mycelial growth, appressorial development, and appressorium turgor but increased conidial production with abnormal morphology [10]; and a gammapartitivirus infection was found to reduce the sporulation of *C. acutatum* [46]. Therefore, most concerns have been attended on the hypovirulent traits for partitiviruses, and no partitivirus-caused hypervirulence was previously reported in the phytopathogenic fungi, except that a partitivirus was found to enhance the virulence of *T. marneffei*, which is a dimorphic clinical fungus causing systemic mycosis in Southeast Asia [15]. As of the current date, there are only two reported cases linking mycoviruses to the hypervirulence of fungi. In one case, a 6.0-kbp dsRNA has been shown to enhance the virulence of *N. radicicola* when horizontally transmitted into fungal strains, while the fungal virulence decreased upon elimination of the dsRNA unit [5]. In the other case, RnQV1 causes significant alterations in pigmentation, rapid growth, and hypervirulence in *L. biglobosa*, as demonstrated by comparisons between virus-infected and virus-cured isogenic fungal strains [6]. Besides, a totivirus was found to increase the production of tenuazonic acid-derived mycotoxinsin in *Magnaporthe oryzae*, which was evaluated by comparison of the secondary metabolite profile of an isolate of APU10-199A (co-infected with a totivirus, a chrysovirus, and a partitivirus) and that of the strain APU10-199A_P lacking the totivirus and chrysovirus [47]. Here, ScPV1 was vertically transmitted to another fungal strain using protoplast transfection with purified virions, the transfectants showed lower growth rates and hypervirulent traits as compared with those of the parent strain, indicating that ScPV1 confers two contradictory effects: growth attenuation and virulent enhancement to *S. camellicola*. Thus, we provide compelling evidence to demonstrate the first instance of a mycovirus, specifically a partitivirus, conferring hypervirulence to a phytopathogenic fungus through protoplast transfection with mycoviral virions. Moreover, most mycoviruses usually confer synergistic effects by attenuating the virulence and reducing the growth to their fungal host, and only Alternaria alternata chrysovirus 1 (AaCV1) was reported to cause two-sided effects to its host fungus (*Alternaria alternata*), which was evaluated by comparison of several virus-high-titer and virus-low-titer isolates of AaCV1-bearing *A. alternate,* in accordance with a 13-fold increase in AK-toxin level [48,49]. To our best knowledge, this is the secondary report that a mycovirus confers tow-sided effect to its fungal host, and this finding represents a significant advancement in understanding the complex interactions between mycoviruses and their host fungi.

In summary, a novel mycovirus named ScPV1, which belongs to the newly proposed genus Epsilonpartitivirus within the family *Partitiviridae*, has been identified and characterized from *S. camellicola* isolated from tea plants. This marks the first report of a partitivirus from a phytopathogenic fungus infecting tea plant. ScPV1 contains two dsRNA segments that encode an RdRp and a CP, respectively, and these segments are encapsidated into isometric virions with divergent sizes. Additionally, the RdRp exhibits an unexpected size and contains some unique amino acids, indicating that ScPV1 possesses unique molecular and morphological characteristics. Biological tests conducted with transfectants generated through protoplast transfection with purified virions rigorously assessed the effects of ScPV1, and demonstrated that ScPV1 impairs the growth of *S. camellicola* while simultaneously enhancing its virulence. This represents the first instance of a mycovirus responsible for the hypervirulence and conferring two-sided effects to a phytopathogenic fungus demonstrated through a virion-transfection manner, as well as the first case of a partitivirus conferring hypervirulence to a phytopathogenic fungus. This study is expected to expand our understanding of viral diversity, evolution, and biological traits of the family *Partitiviridae*.

## Acknowledgements

The authors thank Yan Li and Liyong Zhou from the Agriculture and Rural Affairs Bureau of Zigui County, Yichang City, Hubei Province, for the convenience they provided in the collection of pathogenic fungi infecting tea plants.

## Author Contributions

W.X. conceived this study, designed the investigation and supervised the project. M.C. conducted most of the experiments, and Z.L. wrote the initial manuscript, L.K. conducted some biological test, and W.X., H.A., R.K. and H.M. improved English, presentation, and discussion.

## Data and materials availability

Sequence data supporting the findings of this study has been deposited in GenBank under the accession numbers PQ201539 and PQ201540 for ScPV1. The remaining data are available within the article and the supplementary materials files and from the corresponding author upon request.

